# An improved *Solanum verrucosum* genome provides insight into potato centromeres and epigenetic regulation

**DOI:** 10.1101/2025.10.17.683038

**Authors:** Moray Smith, Amanpreet Kaur, Vikrant Singh, John T. Jones, Ingo Hein

## Abstract

*Solanum verrucosum* is a wild diploid potato species notable for carrying the uncharacterised *Rpi-ver1* gene conferring resistance to *Phytophthora infestans*, possessing unusually structured centromeres, and exhibiting the rare trait of self-compatibility, the genetic basis of which remains unknown. Here, we present a high-quality genome assembly and annotation of *S. verrucosum* accession CPC54. A comprehensive inventory of resistance genes is provided, and candidate genes for *Rpi-ver1* are identified through remapping approaches. The sequence and epigenetic landscape of centromeres are resolved, revealing a mosaic of repeatless and repetitive domains. These centromeres consist of tandem arrays of kilobase-sized repeats, some of which show signatures of transposable element origin. Specific subfamilies of CRM and Tekay elements appear to have adapted to centromeric regions. The basis of self-compatibility in *S. verrucosum* is investigated with respect to the *S-RNase* gene, which is found to be present and intact, but expressed at very low levels. Notably, upstream insertions of transposable elements are present that may interfere with its expression.

## Introduction

*Solanum verrucosum* is a wild diploid potato species native to Mexico. It has been widely studied as it contains valuable sources of disease resistance and can act as a bridge species, enabling crosses between species with different Endosperm Balance Number (EBN) values. In addition, it is self-compatible, unlike most tuberbearing species of Solanum, and has unusual centromeres that are in a mixed state of being repeatless or formed from kilobase-sized repeats (Eijlander et al. 2000; Gong et al. 2012).

Late blight is a serious disease caused by the oomycete pathogen *Phytophthora infestans*, which has impacted potato cultivation and breeding for over 170 years (Ivanov, Ukladov and Golubeva, 2021). A novel, broad-spectrum dominant late blight resistance gene, *Rpi-ver1*, has been identified on chromosome 9 of this species (Z. Liu and Halterman 2006; X. Chen et al. 2018). Resistance to plant viruses and tomato psyllid has also been reported in *S. verrucosum*, while one accession also showed resistance to a broad range of *G. pallida* populations (Carrasco et al. 2000; Castelli et al. 2005; Cooper and Bamberg 2016).

Beyond disease resistance, the unique genomic architecture of *S. verrucosum*, particularly its centromeres, presents an area of research interest. In *Arabidopsis*, centromeres are composed of megabase satellite arrays with 178bp subunits which are organised into higher order repeats, a result of unequal crossover and recombination (Naish et al. 2021). Underpinning this complexity is the cyclic expansion and contraction of ATHILA LTR transposable elements in the centromere, driving evolution and speciation within *Arabidopsis* (Wlodzimierz, Rabanal, et al. 2023). In contrast to *Arabidopsis*, the centromeres of potato and *S. verrucosum* are known to be partially repeatless and, where repetitive, are formed of kilobase-sized repeat subunits (Gong et al. 2012; H. Zhang et al. 2014). These large repeat monomers share sequence similarity with LTR retrotransposons, indicating their role in centromere evolution in *Solanum* (Gong et al. 2012). Another feature of *S. verrucosum* that makes resolving its genome of interest is its self-compatibility and lack of interspecific reproductive barriers (Behling and Douches 2023). This has previously been attributed to the lack of a functional *S-RNase*, although the genetic basis of this remains unknown (Eijlander et al. 2000).

These properties have led to *S. verrucosum* being a focus for multiple genomic studies. For example, in 2019, an assembly of the dihaploid clone 54 was produced through a combination of short-read, long-read, and scaffolding approaches to produce an assembly with a contig N50 of 858kbp (Paajanen et al. 2019). In 2022, an assembly of the monohaploid *S. verrucosum* clone 11H23 was produced with a contig N50 of 21Mbp through PacBio HiFi sequencing and Hi-C scaffolding (A. J. Hosaka, Sanetomo, and K. Hosaka 2022).

Here, we report an improved assembly of the dihaploid *S. verrucosum* accession 54, alongside a comprehensive gene, transposable element, and DNA methylation annotation. Through this assembly, we resolve the complete sequence of the *Rpi-ver1* locus, which contains a single pseudogenised NLR. We also investigate the sequence and epigenetic status of the *S. verrucosum* centromeres, which are revealed to be in a mix of repeatless and repetitive states - the repetitive centromeres are megabase-scaled and formed of large tandem array subunits. Finally, we investigate the status of the *S-RNase* gene in *S. verrucosum*, which is revealed to be densely methylated and likely silenced through transposable element activity in its promoter region.

## Methods

### DNA extraction, sequencing, and assembly

High-molecular-weight (HMW) DNA was extracted from 72-hour dark-treated leaf tissue with the Macherey-Nagel NucleoBond HMW DNA kit following the manufacturer’s specifications.

PacBio sequencing was conducted by the Norwegian Sequencing Centre using the SMRTbell ExpressTemplate Prep Kit 3.0 and sequencing on one 8M SMRT cell on the Sequel II platform using the Sequel II Binding kit 2.2 and Sequencing chemistry v2.0. HiFi reads were generated using the CCS pipeline (SMRT Tools v11.0.0.146107).

For ONT sequencing, HMW DNA was size-selected using a BluePippin 0.75% High Pass cassette with a >15kbp size selection cutoff and libraries prepared with the SQK-LSK109 Ligation Sequencing kit following manufacturer’s specifications. ONT libraries were sequenced on R9.4.1 minION flow cells using 50-75fmol library DNA as input. ONT sequences were basecalled with guppy v4.2.2 using the model dna_r9.4.1_450bps_sup_plant.cfg.

For Hi-C sequencing, libraries were prepared according to the Dovetail® Omni-C® kit protocol. Two independent Hi-C libraries were prepared and sequenced using a NextSeq P2 kit with paired-end 150bp reads.

To assess the homozygosity of the genome, 21-mers were counted and a histogram generated from HiFi reads with kmc v3.2.4 using the options k21 -t10 -m64 -ci1 -cs10000 and kmc_tools transform histogram -cx10000 (Kokot, Długosz, and Deorowicz 2017). The histogram was used as input for genomescope2 v2.0.1 using the default options (Ranallo-Benavidez, Jaron, and Schatz 2020).

A primary genome assembly was produced from the HiFi reads with hifiasm v0.24.0 using the option –primary (Cheng et al. 2021). A secondary assembly was produced from the ONT reads with flye v2.9.5 using the options genomeSize=750m -nanopore (Kolmogorov et al. 2019).

Mitochondrial and chloroplast assemblies were produced from HiFi reads with oatk v1.0 using the option -c 100 and OatkDB v202030921 (Zhou, Brown, et al. 2025). The chloroplast assembly was annotated with GeSeq (Tillich et al. 2017).

To assess genome contiguity and assembly quality, GCI v1.0 was run using the HiFi and ONT reads, which were simultaneously mapped with minimap2 2.28 and winnowmap v2.03 with the default settings (Li 2018; Jain et al. 2020; Q. Chen et al. 2024). BUSCO v3.0.2 was applied to both the genome and protein predictions (Tegenfeldt et al. 2025).

### Hi-C scaffolding

Hi-C reads were aligned to the assembly with BWA v0.7.18 using the options -5SP -T0 and filtered, sorted, deduplicated, and split with pairtools v1.10 using the options –min-mapq 40 –walks-policy 5unique –max-inter-align-gap 3 (Li 2013; Open2C et al. 2023). Alignments were used to scaffold the genome with yahs v1.2a.2 using the option –no-contig-ec (Zhou, McCarthy, and Durbin 2023). The scaffolded genome was prepared for manual curation the Juicer GUI with the yahs juicer pre module and juicertools (Durand et al. 2016). Minor corrections for chromosome fusions were made, and a final assembly was generated.

To label chromosomes, the final assembly was aligned to the DM1-3 516 R44 (v6.1) genome using mashmap v3.0.6 with default settings (Pham et al. 2020; Kille et al. 2023). The alignment was visualised with the D-GENIES web tool, and chromosome identities were inferred (Cabanettes and Klopp 2018).

### RNA-seq analysis

Tissue culture plants of *S. verrucosum* were maintained on MS20 medium and kept in a growth room at a light intensity of 110µmolm^-2^s^-1^, a temperature of 18±2°C, and a photoperiod of 16/8h light/dark. Healthy three-week-old plantlets with fully expanded leaves were selected. *In vitro* shoots with roots were gently removed from the media and dipped for one minute in a zoospore suspension of *P. infestans* isolate W9928C adjusted to 4x10^6^spores/mL. Dip-inoculated microshoots were blotted on sterile paper towels and then planted in fresh MS20 medium in vented containers. The infected plants were kept in darkness for 16 hours and then incubated under the growth conditions mentioned above. The disease severity was recorded by counting the number of leaves showing disease symptoms in 24-hour intervals. The leaf samples from three independent replicates were collected after 0- and 24-hours post infection and immediately immersed in liquid nitrogen before storage at -70°C for further processing.

From each replicate, leaf samples were crushed to a fine powder and 400mg of ground sample was resuspended in 2mL of TRI reagent and vortexed after the addition of 10µL ß-mercaptoethanol. The slurry was left to stand at room temperature for five minutes before centrifugation (10,000g, 10min at 4°C). To the supernatant, 0.2mL chloroform was added (per 1mL) and incubated at room temperature for five minutes before centrifugation (10,000g, 10min at 4°C). The aqueous layer was transferred, 0.5mL isopropanol added, and transferred to a QIA RNAeasy spin column for washing in RPE buffer twice. RNA was eluted in 50µL RNase-free water and the integrity was checked with a Bioanalyzer 2100.

For RNA-seq, samples were checked for a RIN value >8 and processed at the James Hutton Institute’s Genomics facility for generating RNA sequencing libraries using the standard Illumina mRNA Prep kit RNA unique dual UD Indices as recommended, with 100ng total RNA per sample. Libraries were checked on a Qubit fluorimeter and Bioanalyzer 2100 prior to pooling equimolar amounts of individually indexed samples before sequencing. Sequencing was conducted on a NextSeq 2000 at a loading concentration of 750pM using a P3 200 kit, generating paired-end 100bp reads.

RNA-seq mapping and read count estimation was carried out using nf-core/rnaseq v3.12.0, using the default settings. The salmon quantified read counts were imported into R with tximport v3.19 and all subsequent analysis was conducted with deseq2 v3.19. Infection, tissue-specific, and temperature response assays were analysed in separate RNA-seq experiments. Differential expression analysis was conducted with DESeq() using the default settings, followed by lfcShrink() with the apeglm shrinkage estimator. Genes were considered to be differentially expressed if *p*_adj_ *<* 0.01 and |log_2_(FC)*| >* 1.

### Genome annotation

Transposable element annotations were generated with Earl Grey v4.2.4 using the default settings (Baril, Galbraith, and Hayward 2024). To classify LTR elements into clades, libraries were provided to TEsorter v1.4.6 using the rexdb-plant database (R.-G. Zhang et al. 2022).

Tandem repeats were identified with TRASH with the options –win 10000 –m 9000 (Wlodzimierz, Hong, and Henderson 2023). Repeats were classified by their homology to the Earl Grey TE library - a fasta of repeats was extracted from the TRASH .bed file, a non-redundant library was created with seqkit rmdup, and this was used as query against the TE library (Shen et al. 2016). Subject hits with the highest bit score were used to classify repeats according to their LTR clade, and manual validation was carried out to ensure no misclassification occurred. Repeat classifications were merged with the original TRASH .bed file to produce an informative .bed file for use in subsequent analysis.

Gene models were predicted using BRAKER3 v3.0.7 using RNA-seq and the Viridiplantae OrthoDB v.11 database (Kuznetsov et al. 2023; Gabriel et al. 2024). All RNN-seq data were aligned to the masked genome produced by Earl Grey with STAR v2.7.10 (Dobin et al. 2013). To identify NLRs that were not predicted by BRAKER3, Helixer v0.3.0 was applied to the unmasked genome with the model land_plant_v0.3_a_0080.h5 (Holst et al. 2023). Annotations that were identified as NLRs by Resistify and did not overlap with existing BRAKER3 annotations were merged into the final annotation with AGAT.

Resistify v1.1.5 was run with subcommand nlr –retain –coconat to identify complete and partial NLRs with improved CC annotation via CoCoNat, and subcommand prr to identify RLK/RLPs, which uses TMbed to identify transmembrane domains (Bernhofer and Rost 2022; Martin et al. 2022; Madeo et al. 2023; Smith, Jones, and Hein 2025). A phylogenetic tree of NLRs was built from a multiple sequence alignment of NB-ARC domains produced by MAFFT v7.525 with option –auto (Katoh et al. 2002).

Nanopore DNA methylation analysis was carried out using deepsignal-plant v0.1.6 (Ni et al. 2021). Basecalled fast5 files were re-squiggled with tombo v1.5.1 and extracted with deepsignal_plant extract (Stoiber et al. 2017).

### Rpi-ver1 mapping

To identify the *Rpi-ver1* locus, previously generated KASP markers were mapped to the genome with BLASTn and filtered for hits with 100% identity and query length (X. Chen et al. 2018). The *Rpi-ver1* locus was determined to be the locus delimited by the high-confidence KASP markers. To verify that the locus fully represented *Rpi-ver1*, bulk segregant RenSeq reads from the original study were mapped to the *S. verrucosum* genome and filtered with the expected homozygosity for the parent datasets, and heterozygosity for the F1 progeny bulks.

### Centromere analysis

Existing CENH3 ChIP reads for *S. verrucosum* (SRR18548893) were aligned to the genome with bowtie2 v2.5.3 (Langmead and Salzberg 2012). Centromere coordinates were determined manually by inspecting each chromosome for CENH3 read peaks. To identify transposable elements with a bias for presence in the centromere, Fisher’s exact tests were conducted for each family identified by Earl Grey, using inside/outside centromere counts. Only intact transposable element annotations were counted, and only families that could be additionally identified by TEsorter were investigated further. A phylogenetic tree of reverse transcriptase domains was built using iqtree -bb 1000 -nt AUTO from a domain alignment produced by TEsorter.

### Characterisation of the S-locus

To identify the homolog of *S-RNase* in *S. verrucosum* 54, known *S-RNase* sequences from *S. neorickii* and *S. chilense* (BAC00940.1 and BAC00934.1) were queried against *S. verrucosum* with BLASTp v2.9.0. The *S-RNase* homolog was determined by its higher percentage identity relative to non-S-locus RNases and its expected location on chromosome 1. The gene model predicted by BRAKER3 was cross-validated against the Helixer annotation and found to be identical.

The conserved sequence was used to design *S-RNase* specific primers (fwd: CTGGCCTCAACT CAGATACGA, rev: ATTCCATGATTTCTAAGAGTTCCC) for gene quantification. RNA was extracted from flowers of *S. verrucosum* 54 using the previously described methodology, and cDNA was synthesised using the SuperScript IV first strand cDNA synthesis kit (Invitrogen) following the manufacturer’s guidelines. The housekeeping gene exocyst complex component (*Sec3*) was amplified using primers (fwd: GCTTGCACACGCCATATCAAT, rev: TGGATTTTACCACCT TCCGCA) for gene quantification. Relative gene expression was calculated using the formula ΔCt = Ct_SRNase_ − Ct_Sec3_.

### Data analysis

Where not otherwise specified, analysis was conducted using R v4.5.0 - scripts and packages were controlled by renv and are available at https://www.github.com/swiftseal/sver54_assembly. The bioinformatics and computational analyses were performed on Crop Diversity HPC, described by (Percival-Alwyn et al. 2024).

## Main

### A comprehensive genome assembly of *S. verrucosum*

To generate a high-quality assembly of *S. verrucosum*, a combination of long-read sequencing methods was selected. A total of 39.5Gbp of HiFi reads with an N50 of 14.8kbp and an average quality of 28.7 were generated, producing a total assembly size of 773.8Mbp across 2,129 contigs with a contig N50 of 46.3Mbp. Simultaneously, a total of 39.7Gbp of ONT reads were generated with an N50 of 14.3kbp and an average quality of 13.63, producing a total assembly size of 686.4Mbp across 1651 contigs with an N50 of 5.3Mbp. As the HiFi assembly produced an assembly with a 10-fold higher contiguity, this was taken forward for scaffolding with Hi-C sequencing. A total of 502 million read pairs were generated, of which 293 million were mapped, and 21 million were marked as non-duplicate and valid contact events. Scaffolding from these produced a 696Mbp chromosome-scale assembly composed of 19 contigs, 12 scaffolds, and a contig N50 of 46.3Mbp (Figure 1a).

**Figure 1:**
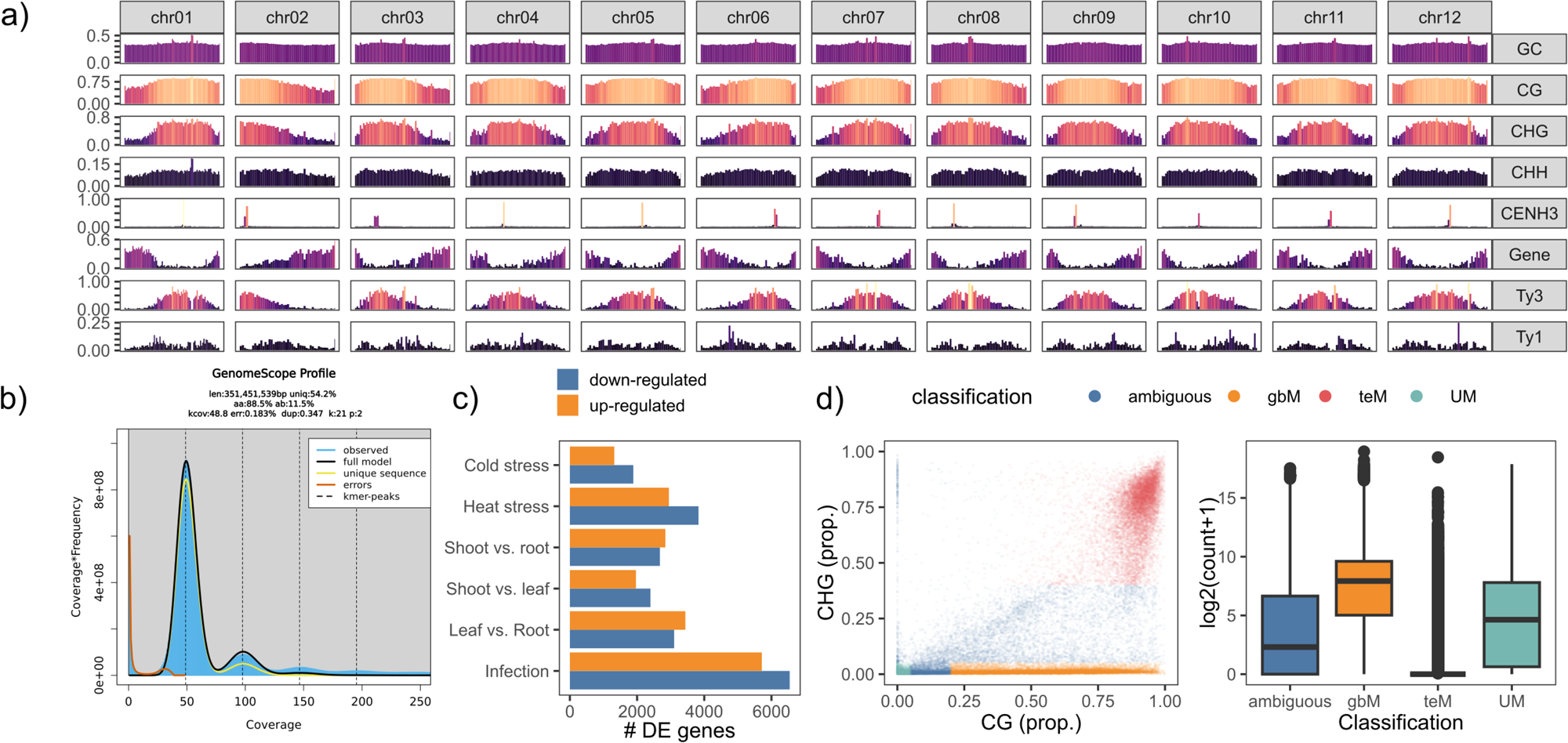
a) The density of GC%, CG, CHG, and CHG methylation, CENH3 read mapping, genes, and Ty1 and Ty3 LTRs across the genome assembly. b) The GenomeScope2 profile of 21-mers extracted from HiFi reads. c) The number of differentially expressed genes identified across RNAseq conditions. d) Left: The distribution of genes by their CG and CHG exon methylation, coloured by their classification. Right: the leaf RNA expression of different methylation classifications.

Many of the assembled chromosomes could be described as “telomere-to-telomere”; however, this claim can be over-inflated by assembly artifacts and misassemblies (Q. Chen et al. 2024). To further assess the quality of the final genome assembly, the Genome Continuity Inspector (GCI) was used to evaluate contiguity and identify potential misassemblies. The complete assembly had a GCI score of 60.8, with 6/12 chromosomes being represented by a single contig, whilst the remaining were limited to two to three contigs (Supplementary Table 1). Only a single locus on chromosome 9 was noted for a significant drop in read coverage across both HiFi and ONT reads, indicating a potential misassembly (Supplementary Figure 1).

As expected, the genome is diploid and 88.5% homozygous according to GenomeScope2 (Figure 1c). A high BUSCO completeness (C: 99.1% [S: 83.8%, D: 15.3%], F: 0.7%, M: 0.2%, n: 425) also demonstrated a high-quality assembly. Overall, this assembly represents a marked improvement over the previous best *S. verrucosum* assembly, which had a contig N50 of 21Mbp (A. J. Hosaka, Sanetomo, and K. Hosaka 2022).

For improved gene annotation, RNA-seq was conducted across a variety of tissue levels (root, shoot, and leaf), biotic (infected with *P. infestans*) and abiotic stress conditions (heat and cold) (Supplementary Table 2). Genes were annotated with a combination of BRAKER3 and Helixer, identifying 38,710 genes with a mean gene length of 3043bp, 1.2 transcripts per gene, 4.3 exons per transcript, and a multi-to-single exon ratio of 2.75. The BUSCO completeness of the annotation was also high (C: 99.8% [S: 99.8%, D: 0.0%], F: 0.2%, M: 0.0%, n: 425). RNA-seq analysis 24 hours post infection of *P. infestans* isolate W9928C revealed higher gene down-regulation as compared to up-regulation. Similarly, during cold and heat stress, more genes were down-regulated. Differential expression of genes was also observed to be organ-specific (Figure 1c).

Two circular mitochondrial genomes were assembled, which were 417.9kbp and 49.3kbp in size, respectively. A single circular chloroplast genome of 155.5kbp was also assembled (Supplementary Figure 2). This was in line with a recent 155.5kbp assembly of the *S. verrucosum* chloroplast which was contrastingly lacking in a ycf1 annotation at the IRB-SSC boundary (L. Zhang et al. 2024).

DNA methylation is an important characteristic of gene and transposable function, and so DNA methylation was determined from the available ONT reads. The average level of methylation in the genome was 70%, 46.2%, and 9.4% in the CG, CHG, and CHH contexts, respectively - in line with previous estimates for *S. lycopersicum* and *S. melogena* (Cui et al. 2021; Lu et al. 2021). A clear trend of heightened methylation towards the centromeres was evident for all chromosomes, across all methylation contexts (Figure 1a).

In plants, genes can be classified according to the levels of CG and CHG methylation in their exon body (Zeng, Dawe, and Gent 2023). Accordingly, they can be classified as unmethylated (UM) if unmethylated in either context, or as having gene body methylation (gbM) if methylated in the CG context only, or as having TE-like methylation (teM) if methylated in both the CG and CHG contexts. In maize, it has been previously observed that gbM are generally highly expressed across all tissues, that UM genes are more frequently tissue-specific, and that teM genes are transcriptionally silent (Zeng, Dawe, and Gent 2023). To determine if a similar effect is observed in *Solanum*, the same classification criteria were applied to the *S. verrucosum* genome.

Accordingly, 5,940 genes were classified as UM, 17,151 as gbM, 8,076 as teM, and 7,957 as ambiguous (Figure 1d). Although the maize genome has a similar number of genes (39,000), more genes were classified as being gbM (17,151 vs 8,134) and teM (8,076 vs 3,402) in *S. verrucosum*. In line with the previous investigation, genes classified as teM had substantially lower expression than those classified as gbM (Figure 1d). No evidence of a tissue-specific expression bias was found for UM genes, likely because far fewer tissue-specific expression datasets were available for this study.

To create a high-quality transposable element annotation, the performance of TE annotation tools EDTA and Earl Grey were compared (Supplementary Figure 3). Earl Grey classified 67.6% of the genome as being repetitive, whereas EDTA classified 56.7%. As expected, the largest proportion of repeats was classified as being LTR-derived, with both pipelines identifying similar proportions of 39.7% and 36.6%. Earl Grey identified significantly fewer LTR families (n=632) than EDTA (n=1975), which were also of greater mean length. A similar trend was also seen for DNA, Helitron, and SINE elements, but not LINEs, for which Earl Grey identified a larger number of families.

To further assess the completeness of the TE libraries assembled by both tools, TEsorter was applied to classify TEs by their domains. The fraction of LTRs that failed to be classified by TEsorter was 63.1% and 9.8% for EDTA and Earl Grey respectively. TEsorter further classified 38.6% (n=252) of LTRs as being complete in the Earl Grey library, but only 15.5% (n=306) in the EDTA library.

The smaller number of TE families, greater mean length of LTR families, and higher rate of successful LTR classification by TEsorter were taken as evidence of Earl Grey outperforming EDTA at producing a high-quality TE library. Therefore, all subsequent analysis was undertaken using the Earl Grey library.

In the genetic history of *S. verrucosum*, two potential bursts of LTR insertion are noted, indicated by the two peaks in the repeat landscape (Supplementary Figure 4). The most recent burst also appears to have been associated with DNA type transposable elements. Interestingly, upon closer examination of the LTR history it appears that the recent activity was associated with Tekay, Athila, and CRM Ty3 LTRs, whereas the Ty1 Clades TAR, Ikeros, and Bianca contributed to a peak of activity approximately between the two transposable element bursts.

Together, the curation of a high-quality assembly, DNA methylation dataset, and gene and transposable element annotations provides a valuable resource for investigations into the *S. verrucosum* genome. Below, studies into the genomic basis of disease resistance, centromere organisation, and self-compatibility are described.

### Taking inventory of resistance genes

Nucleotide-binding leucine-rich repeat genes (NLRs) frequently function as the sensors of pathogen-derived signals or their function, and as such are a vital component of the plant’s capacity to elicit an immune response to disease. In *S. verrucosum*, a total of 490 NLRs were identified (Figure 2a). Of these, 238 were classified as canonical coiled-coil NLRs (CNLs), 44 as toll-interleukin receptor NLRs (TNLs), and two as RPW8 NLRs (RNLs). Of the TNLs, 28 had C-JID domains, and 41 of the CNLs had MADA motifs (Figure 2b). The majority of LRR domains were 350aa in length, although several NLRs with LRRs longer than 1,000aa were noted (Figure 2c).

**Figure 2:**
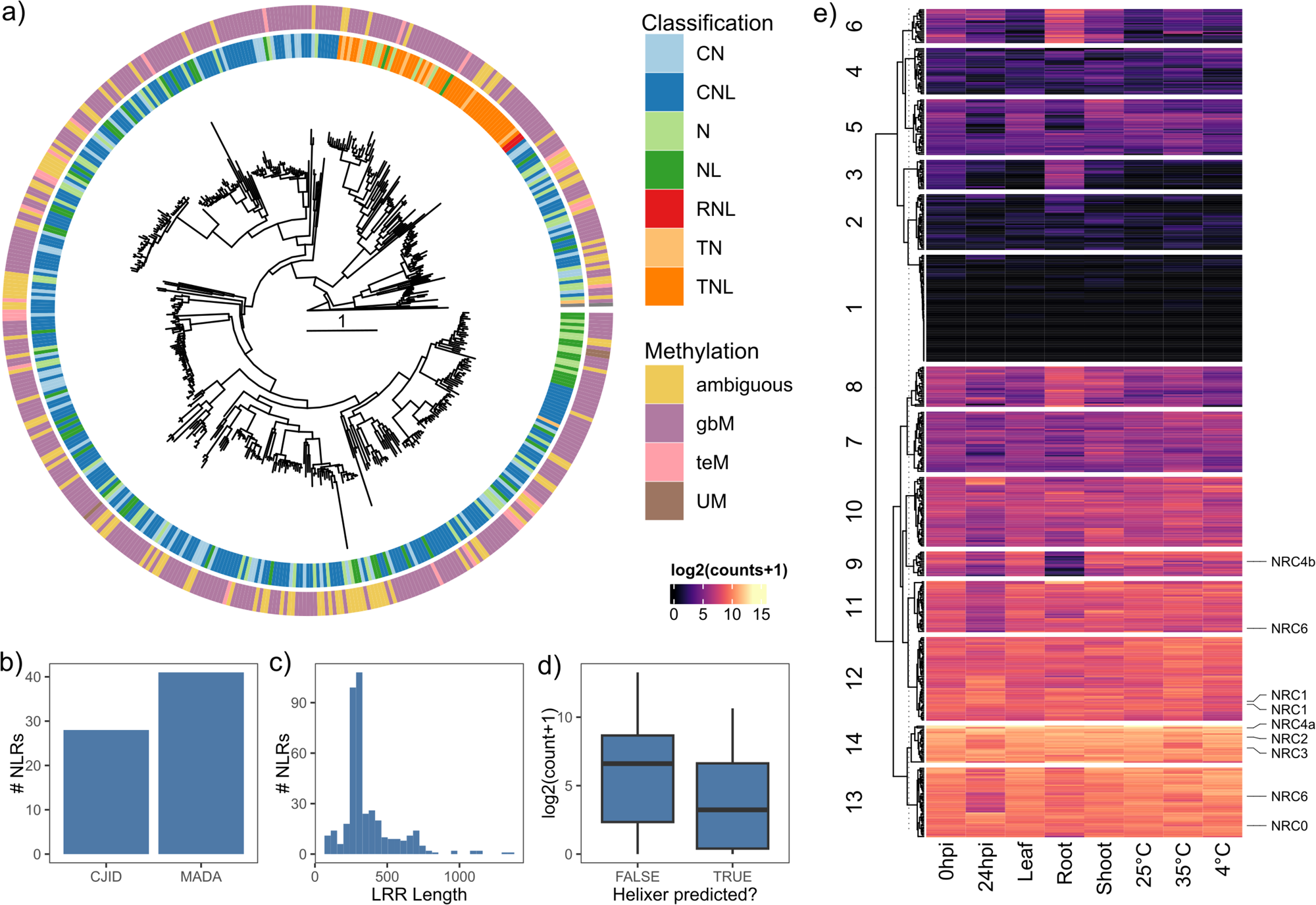
a) Phylogenetic tree of NLR NB-ARC domains identified in this study, rooted to CED-4 from C. elegans. b) Number of NLRs with C-JID and MADA motifs identified. c) Histogram of LRR lengths. d) Leaf expression of NLRs identified by BRAKER3 or by Helixer. e) Clustered heatmap of NLR expression across conditions explored in this study. NRCs have been highlighted.

In the process of annotating the genome, it was noted that the gene predictor Helixer identified substantially more NLRs than BRAKER3 alone. In total, 163 NLRs were identified solely by Helixer, indicating that a large fraction of NLRs evade BRAKER3 annotation. Helixer-identified NLRs had significantly lower expression (ß=-1.97 log2(count+1), SE=0.33, p=7.55e-09), suggesting that they could function as valid NLRs, and that their evasion of BRAKER3 is possibly expression-led (Figure 2d).

Indeed, of the identified NLRs, 23.1% exhibited low to no expression across all RNA-seq conditions explored in this study (Figure 2e). When NLRs were clustered based on their expression patterns across RNA-seq conditions, root tissue emerged as a clear outlier for a subset of NLRs, consistent with previous findings (Dahlen et al. 2023). However, previously, it has been reported that a subset of NLRs - including *Hero* and *NRC6* - have root-specific expression in *Solanum* (Lüdke et al. 2023). Whilst two *NRC6* homologs exist in *S. verrucosum* and root-specific NLR clusters were visible, no NRCs exhibited root-specific expression and instead spanned three separate clusters of varying NLR expression. It should also be noted that in the previous study, only one copy of NRC6 was identified. The second copy identified in this study was identified solely through Helixer, demonstrating the importance of capturing a complete NLR inventory prior to downstream analysis.

In addition to the NLRs, a total of 705 RLKs and 517 RLPs were identified. Of these, extracellular LRR domains were the most common (35%, n=423), followed by G-LecRLK (11%, n=131) and WAK (6%, n=71) domains. An additional 431 RLK/RLPs had extracellular domains that did not fit into the classification schema outlined in the RLKdb (Z, J, and D 2024).

Previous mapping experiments in *S. verrucosum* isolated the novel disease resistance gene *Rpi-ver1* to a 4.3Mbp locus on chromosome 9 in the DM reference genome. Re-analysis of existing mapping data - in particular of the markers DMG400017237 and DMG400017146, which demark this locus - reduced this to a locus 1.4Mbp in the new genome from 54.6Mbp to 56.0Mbp. Within the locus are two adjacent, NLR-like genes that appear to be a canonical CNL with a frameshift mutation in the NB-ARC domain. In addition, there is an Rx-CC Jacalin-like lectin domain protein and a cysteine-rich kinase RLP, both of which have been previously implicated in disease resistance in plants (Esch and Schaffrath 2017; Kamel et al. 2023).

### The sequence and epigenetic status of centromeres

To identify putative centromeric regions, previously generated CENH3 ChIP reads were realigned to the *S. verrucosum* genome (H. Zhang et al. 2014). In total, 98% of the reads aligned to the primary chromosome assembly, indicating that the vast majority of centromere sequence had been successfully assembled, an improvement on a previous assembly where 17.2% of reads mapped to unanchored contigs (A. J. Hosaka, Sanetomo, and K. Hosaka 2022). Clear peaks of CENH3 ChIP read mapping were present on each chromosome (Figure 1a) and accordingly, centromere regions were identified (Figure 3a). Given the dynamic nature of centromeres, it should be noted that the CENH3 ChIP reads used in this study were from a previous study of a different *S. verrucosum* line - as a result CENH3 mapping peaks are unlikely to be fully representative of the true state of CENH3 mapping in this genome (H. Zhang et al. 2014).

**Figure 3:**
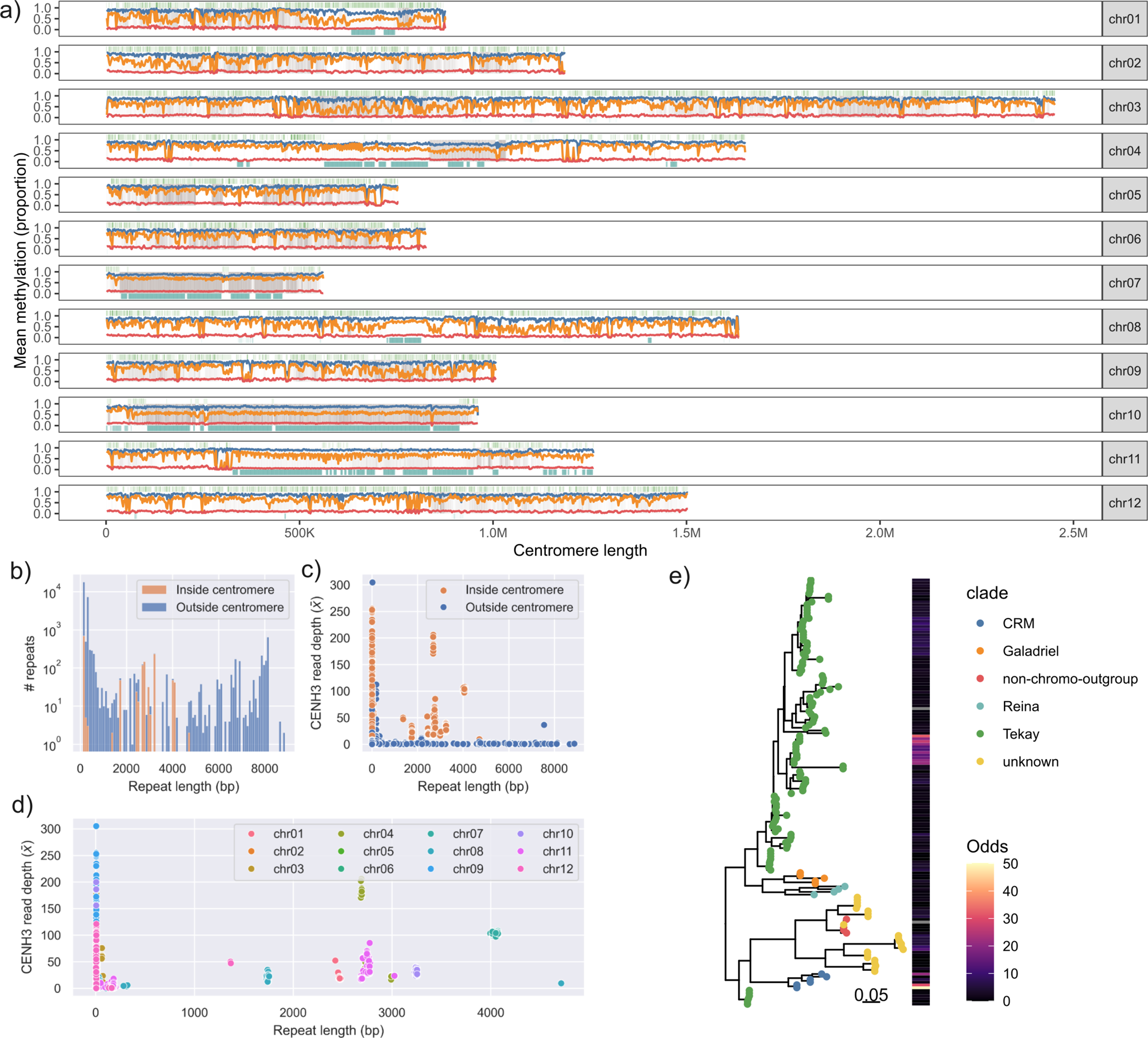
a) The density of LTRs (top, green), CENH3 read mapping (middle, grey), TRASH-derived repeats (bottom, teal), and CG, CHG, and CHG methylation (blue, orange, red, respectively) across *S. verrucosum* centromeres. b) Histogram of length of repeats identified across the genome. c) Distribution of repeats by length and average CENH3 read depth. d) Distribution of centromeric repeats by length and average CENH3 read depth, coloured by centromere identity. e) Subset of phylogenetic tree of LTRs identified in this study built from the RT domain. LTRs coloured according to their clade identified by TEsorter. The odds ratio of centromere bias from Fisher’s exact test are highlighted.

Previous studies have shown that the centromeres of potato and *S. verrucosum* are either repeatless or formed of tandem repeat arrays with unusually large kilobase-sized subunits. As conventional transposable element annotators often fail to reliably detect tandem repeats, TRASH was applied to identify tandem repeat arrays with subunits less than 10,000kbp. This analysis revealed a diverse range of large repeat subunits across the genome, including within the centromeric regions (Supplementary Table 3). As expected, only a subset of centromeres were repetitive, specifically cen04, cen07, cen08, cen10, and cen10, whilst the remainder appeared repeatless. Notably, cen01, cen08, and cen11 showed a mixture of repeatless and repetitive sequences enriched for CENH3 binding. Although repeat subunit sizes varied among centromeres, all were within the kilobase range. Most centromeres consisted of a uniform tandem array, except cen04, which contained multiple distinct arrays. In line with previous observations, centromeric repeats were enriched for CENH3 mapping relative to non-centromeric repeats (Figure 3c). Interestingly, repeats for each centromere appeared to be unique in size (Figure 3d).

Manual inspection of the centromeric regions also indicated the presence of intact transposable element insertions, including insertions that disrupted the repetitive arrays. To identify transposable element families with a disposition for centromere presence, their bias towards presence in the centromere was calculated. From this, it appears that the CRM LTRs and a subgroup of Tekay LTRs have specialised to invade the *S. verrucosum* centromeres (Figure 3d).

Centromeres are dynamic structures, and previous studies have indicated that the centromeres of *S. verrucosum* and *S. tuberosum* are divergent (H. Zhang et al. 2014). As an example, a comparison of the sequence of cen7 revealed little sequence similarity between *S. verrucosum* and *S. tuberosum* (Supplementary Figure 5). In line with the observation that *S. verrucosum* centromeric arrays are mostly uniform for each chromosome; no higher-order repeat topology was observed in either centromere.

In the centromeres of *Arabidopsis* and pepper, methylation in the CHG context is relatively reduced, potentially due to depletion of H3K9me2, which maintains methylation homeostasis (Naish et al. 2021; K. Zhang et al. 2025). A similar trend was not observed for *S. verrucosum*, where no clear correlation between centromeric repeats is observed, beyond the general trend of increased methylation towards the centromeric locus (Figure 3a).

The repeatless centromeres of *S. verrucosum* are similar in structure to two recent assemblies of the *C. annuum* and *C. rhomboideum* genomes (W. Chen et al. 2024). In both cases, the centromeres are formed of a rich landscape of Ty3 LTRs, with members of the Athila, Tekay, and CRM clades being present. In *Capsicum*, CRM LTRs are particularly dominant in the repeatless centromeres. One explanation for this enrichment is that CRM clades have specialised chromodomain and CR motifs which enable centromere targeting through an unknown mechanism (Neumann et al. 2011). The absence of these structural elements in families such as the centromere-biased Tekay LTRs observed in *S. verrucosum* indicates that other mechanisms exist which drive centromere insertion or retention. Recently, a mechanism that drives centromere-bias of LTRs has been elucidated for the ATHILA family in *Arabidopsis*, whereby centromeres become “addicted” to ATHILA insertions due to them silencing transcription whilst simultaneously providing small RNAs that restore normal centromere function (Shimada et al. 2023).

### The status of *S-RNase*

The self-compatibility of *S. verrucosum* has been attributed to a lack of a functional *S-RNase* in the genome, either through mutation, a lack of expression, or complete absence. A search for the *S. neorickii S-RNase* (BAC00940.1) revealed a single high-scoring hit cpc54_3362 (Identity: 48.9% E-value: 9.78e^-68^) on chromosome 1, the expected location of the S-locus.

The sequence of the *S. verrucosum S-RNase* gene was examined for any mutations that could result in non-functionality. No evidence of any truncations or significant insertions that might lead to a loss of function was observed. Previously identified mutations that might have led to reduced or a loss of *S-RNase* function including a lost N-glycosylation site and histidine active sites are also not present (Broz et al. 2021), suggesting that the *S-RNase* of *S. verrucosum* is likely to be functional.

No expression of the gene cpc54_3362 was identified in any of the RNA expression analyses. However, since the RNA-seq analysis did not specifically include flower tissue, we conducted a qRT-PCR analysis to assess *S-RNase* expression in this tissue. In flowers of *S. verrucosum* 54, *S-RNase* expression was detected with a Ct value of 32.6, relative to the housekeeping gene *Sec3* (Ct 26.6), corresponding to a relative expression level of 0.0156. Therefore, although *S-RNase* was expressed, this very low expression level may contribute to the self-compatibility observed in this accession.

The *S-RNase* gene is located in a pericentromeric region that is exceptionally dense in LTRs and other transposable elements. Analysis of the 2kbp upstream region also revealed a high density of TEs (Figure 4). However, annotations produced by Earl Grey and EDTA differed. EDTA identified two TIR fragments belonging to the Mariner and CACTA families within a repetitive region, whereas Earl Grey did not classify these elements fully. Instead, Earl Grey identified two large LINE/L1 fragments and an intermediate MULE-MuDR element in the same region.

**Figure 4:**
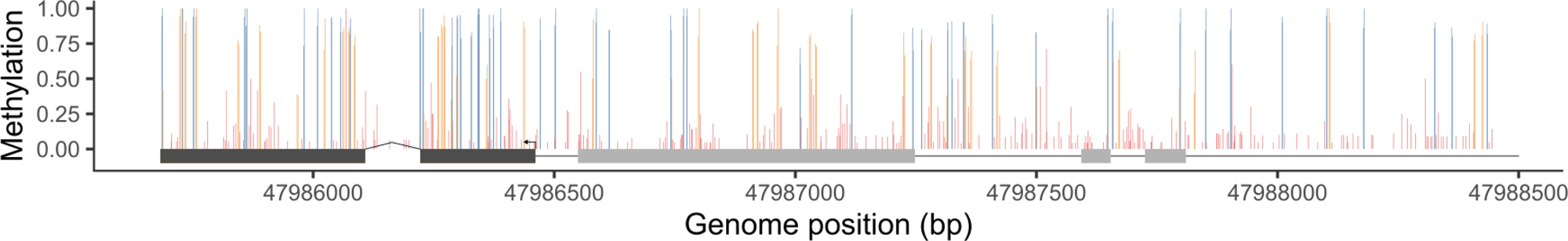
The epigenetic state of *S-RNase* in *S. verrucosum*. The *S-RNase* exon sequence is highlighted in dark grey and identified upstream TIR tranposable element sequences in light grey. The proportion of methylation in the CG (blue), CHG (orange), and CHH (red) contexts is also visualised.

Transposable element activity at the *S-RNase* locus is common to angiosperms, likely driving their diversification through relocating or shuffling the gene structure (Y. Liu et al. 2025). In citrus, MITE transposable element insertions have directly led to self-compatibility transitions and are indirectly associated with tomato self-compatibility alleles (Broz et al. 2021; Hu, C. Liu, et al. 2024).

The self-incompatible *Nicotiana alata* has a Ty3 LTR insertion upstream of its *S-RNase* and whilst it does not impact expression directly, it may be the causal element behind enrichment of CHH methylation near the locus in the pistil, where it is expressed in abundance (Salcedo-Sánchez, Cruz-Zamora, and Cruz-García 2023). Recently, a novel mechanism of self-compatibility has been observed in *Poncirus trifoliata* whereby recombination has resulted in the formation of a “super S-haplotype” containing two copies of the S-locus from independent haplotypes (Hu, C. Liu, et al. 2024). The two S-locus copies cause self-recognition in the pollen, breaking self-incompatibility and leading to self-compatibility. This rare recombination event is posited to be driven in part by MITE insertions, owing to their presence at the recombination breakpoint.

Likely as a result of transposable element activity, the *S-RNase* gene and its surrounding locus in *S. verrucosum* are densely methylated in the CG and CHG contexts, but not in the CHH context. This methylation pattern is likely to affect gene expression. In almond, DNA methylation of the *S-RNase* promoter region has been closely linked to the transition to self-compatibility (Fernández i Martí, Gradziel, and Socias i Company 2014). In *Nicotiana alata*, the promoter region of *S-RNase* accumulates dense CHH methylation specifically in pistil tissue, where the gene is highly expressed (Salcedo-Sánchez, Cruz-Zamora, and Cruz-García 2023).

Interestingly, a recent study of a self-compatible citrus mutant reported no change in average DNA methylation across the *S-RNase* locus in style tissue. However, a distinct CHH methylation island was identified upstream of the locus, present exclusively in the self-compatible mutant (Hu, Xu, et al. 2021). These findings together suggest that both the presence and context of DNA methylation in the *S-RNase* promoter region may play key roles in regulating expression and influencing self-compatibility in *S. verrucosum*.

## Conclusions

Here, a high-quality assembly of the *S. verrucosum* genome is presented alongside gene, transposable element, and DNA methylation annotations.

The full complement of NLRs and RLK/RLPs is resolved for *S. verrucosum*, highlighting that a significant proportion of expressed NLRs evade identification by standard gene annotation approaches such as BRAKER3. Re-analysis of existing *Rpi-ver1* mapping data reduced the size of the resistance locus to a locus on chromosome 9 between 54.6Mbp and 56.0Mbp and revealed its full sequence identity. Key candidates of the *Rpi-ver1* resistance gene include a pseudogenised NLR, a Rx-CC Jacalin-like lectin domain protein, and a cysteine-rich kinase RLP. Resolving the *Rpi-ver1* locus enables the design of markers for future fine-mapping experiments to identify the underlying gene.

The complete sequence of the *S. verrucosum* centromeres is also presented, revealing a complex of repeatless and repetitive sequences that appears to be actively shaped by transposable elements. Given the relative ease with which the *S. verrucosum* genome is assembled, we anticipate that future studies on the trans-generational dynamics of its centromeres will be valuable in elucidating their plasticity. An attractive hypothesis is that the repeatless centromeres may function as transitory centromeric structures until a repetitive structure can be seeded and expanded. DNA methylation is not a predictive mark of *S. verrucosum* centromeres - the identification of other centromere-specific marks, particularly chromatin modifications, will be beneficial in identifying epigenetic features that define the repetitive and repeatless centromeres.

Finally, the presence of an intact *S-RNase* gene in *S. verrucosum* was confirmed. The gene shows no structural abnormalities that would suggest non-functionality, but it is lowly expressed and densely methylated, likely due to transposable element activity in its promoter region.

## Declarations

### Ethics approval and consent to participate

Not applicable.

### Consent for publication

Not applicable.

## Availability of data and materials

All analysis code is available at https://github.com/swiftseal/sver54_assembly. Genome and annotation data is available at https://doi.org/10.5281/zenodo.17107540. Read data is available under PRJEB97628 ERP180128.

## Author Contributions

MS conducted all data analysis presented in this manuscript. AK conducted all RNA-seq tissue and RNA preparation. VS prepared and analysed qPCR expression data. MS, JJ, and IH contributed to the planning of the study. All authors substantively revised the manuscript. All authors read and approved the final manuscript.

## Acknowledgements

The authors acknowledge the Research/Scientific Computing teams at The James Hutton Institute and NIAB for providing computational resources and technical support for the “UK’s Crop Diversity Bioinformatics HPC” (BBSRC grant BB/S019669/1), use of which has contributed to the results reported within this paper.

The PacBio sequencing service was provided by the Norwegian Sequencing Centre (https://www.sequencing.uio.no), a national technology platform hosted by the University of Oslo and supported by the “Functional genomics” and “Infrastructure” programs of the Research Council of Norway and the Southeastern Regional Health Authorities.

---

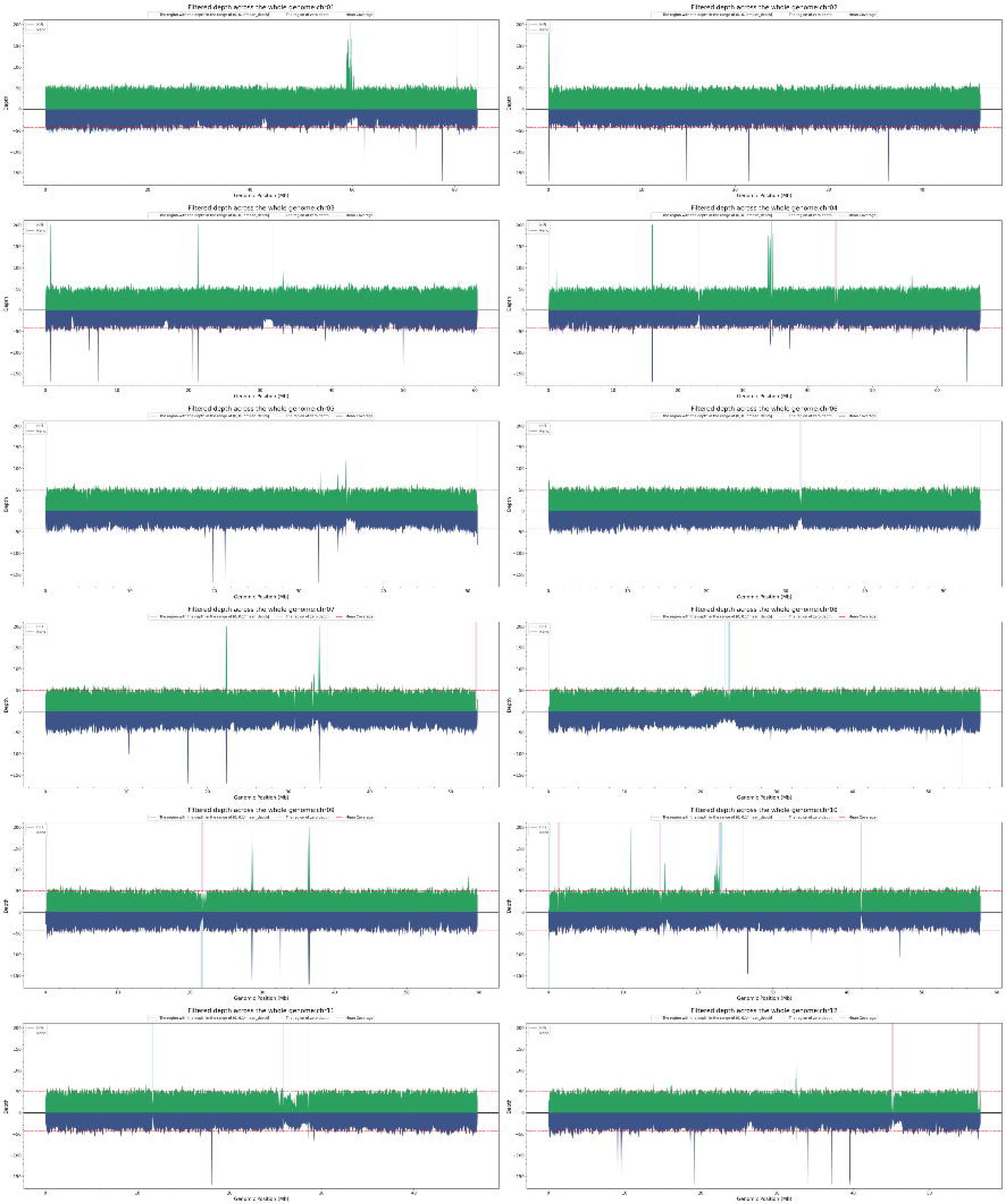

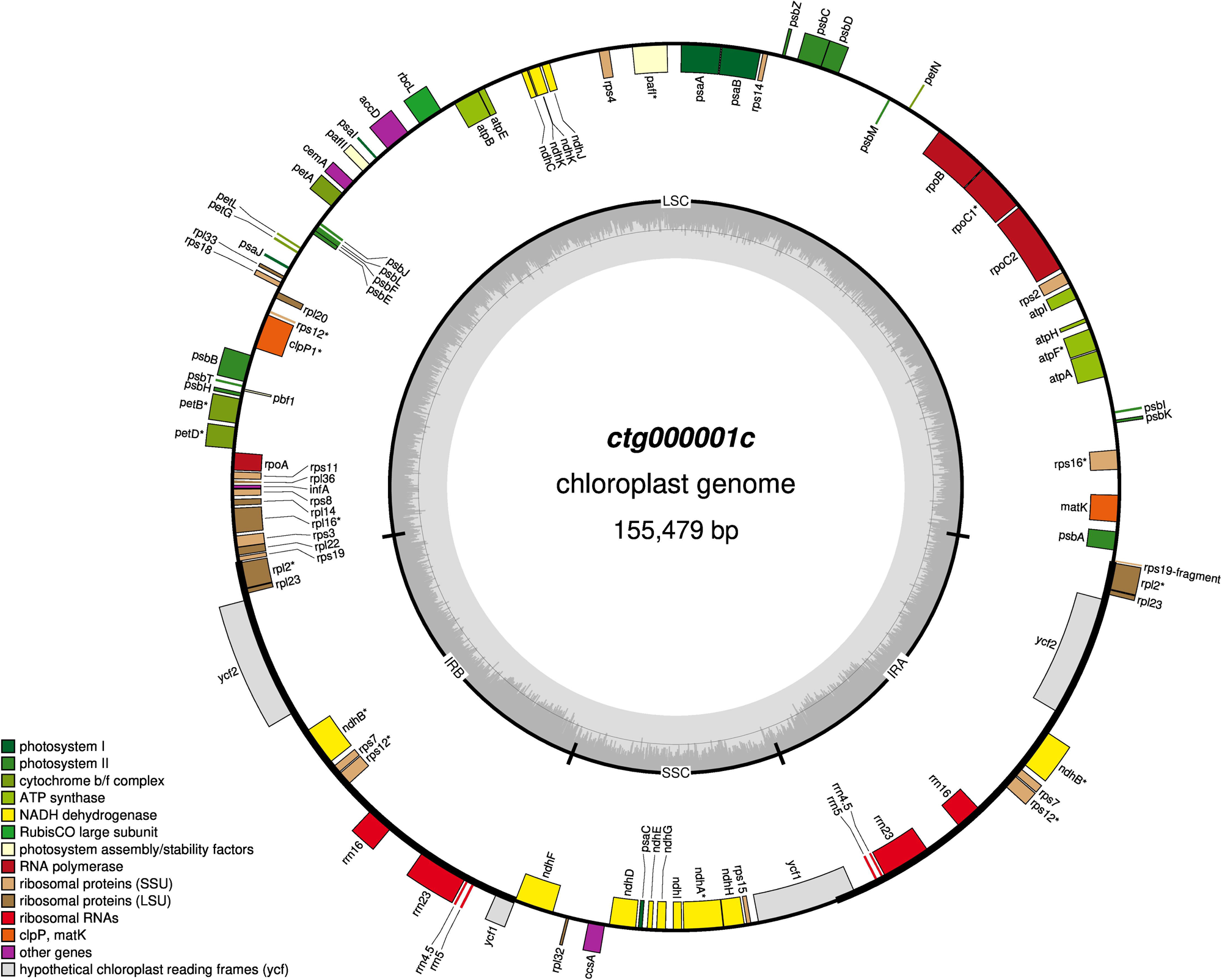

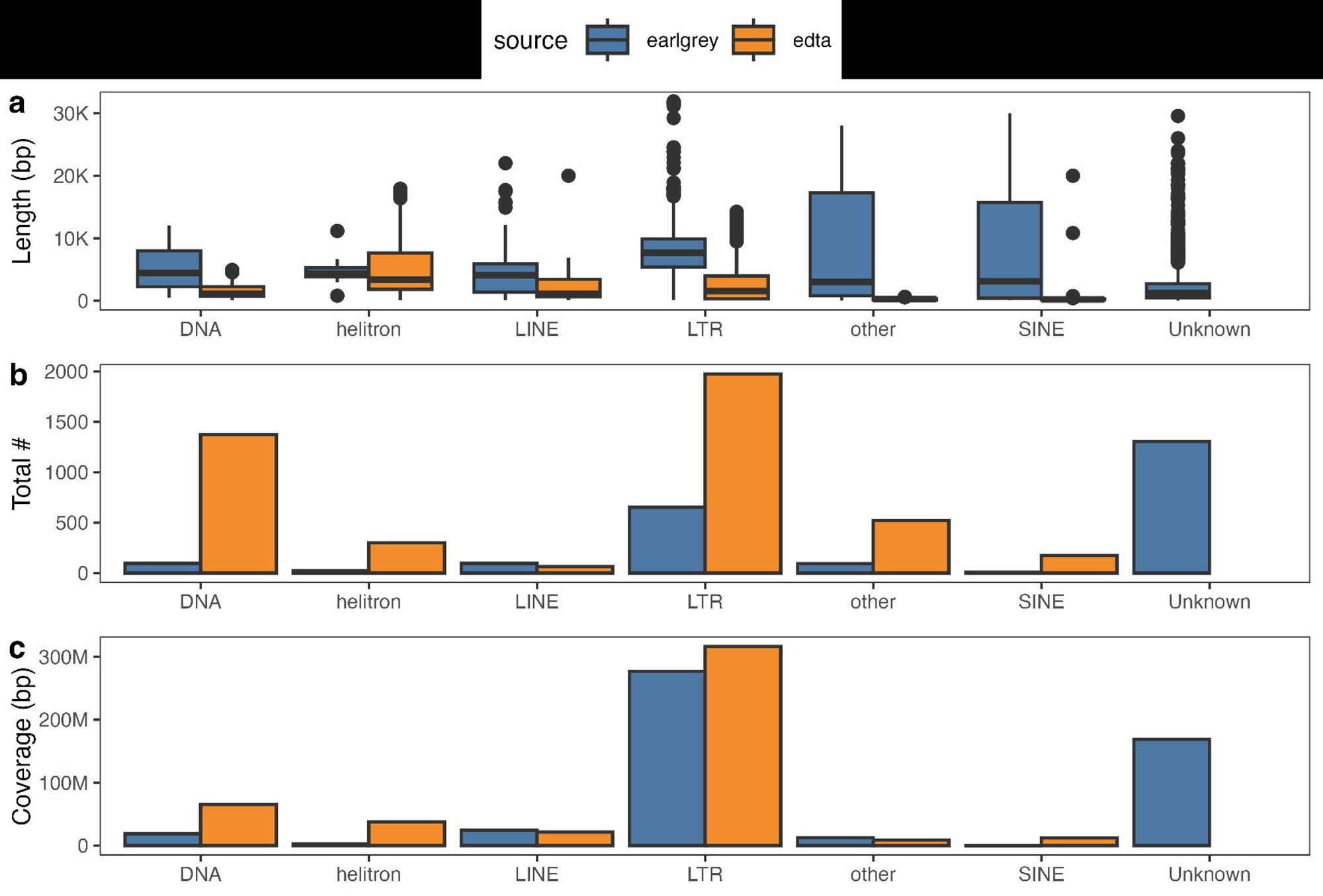

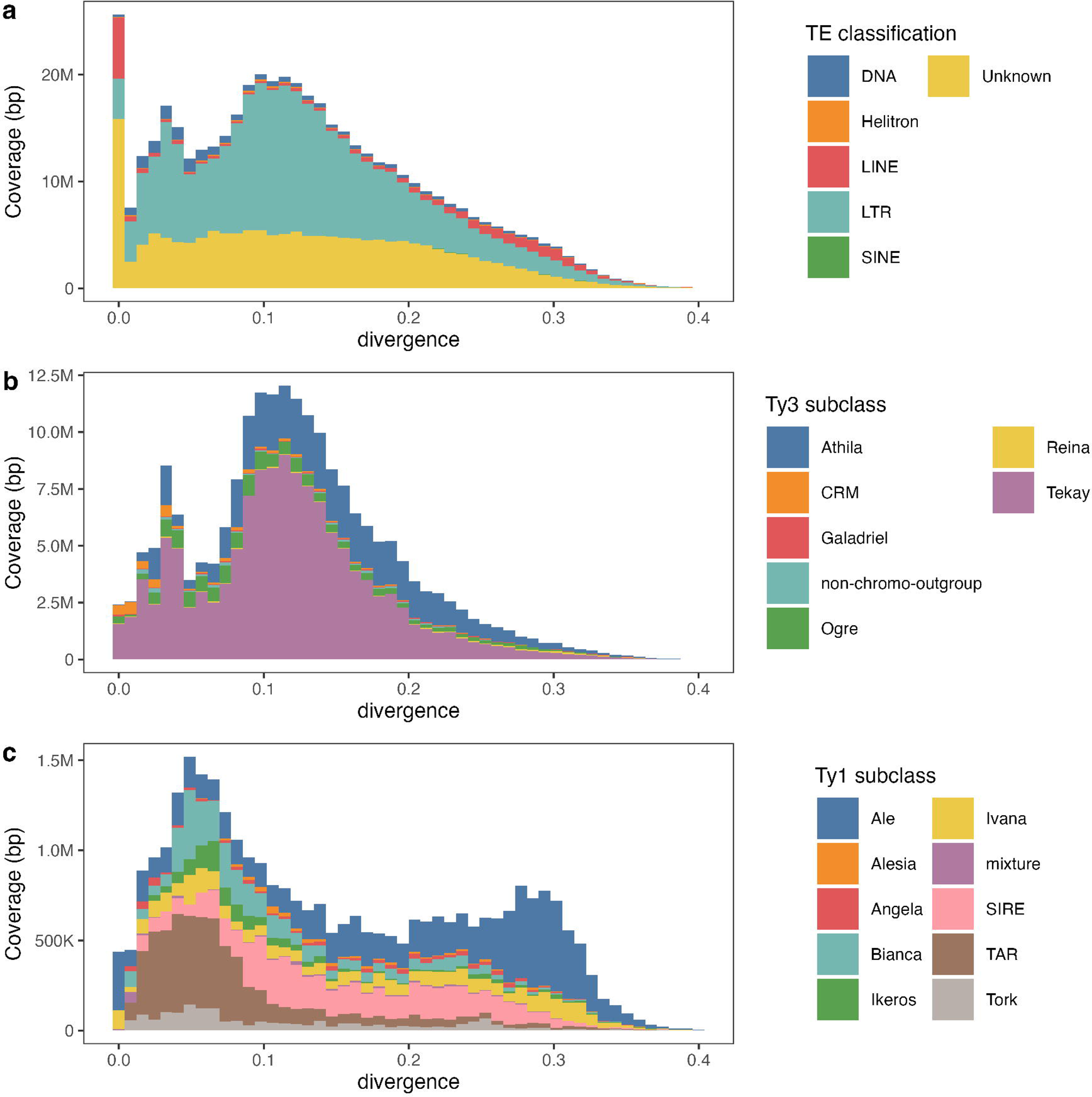

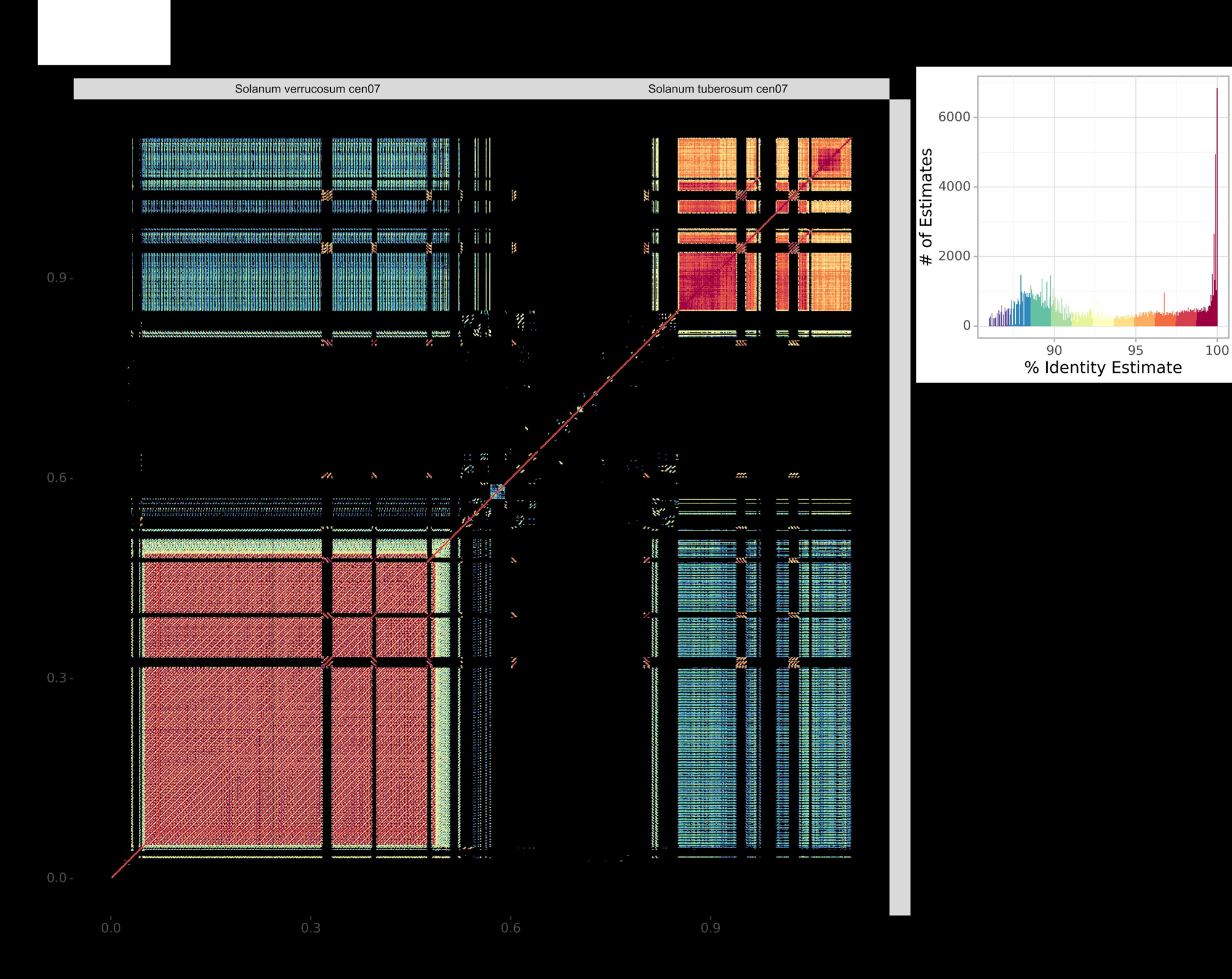

